# Divergent RNA structures support accurate splicing of the SF3B1-sensitive *MAP3K7* intron

**DOI:** 10.64898/2026.05.19.726294

**Authors:** Austin Herbert, Alexandra Randazza, Abigail Hatfield, Lela Lackey

## Abstract

Splicing is governed by interactions between the spliceosome and precursor RNA sequence and structural elements. However, the relative contributions of RNA sequence and structural elements remain unclear. Here, we systematically dissect these determinants using a high-throughput mutagenesis approach with the *MAP3K7* intron reporter. The *MAP3K7* gene encodes a serine/threonine kinase involved in response to environmental stress. *MAP3K7* precursor RNA contains a cryptic 3’ splice site that increases in use when the core spliceosomal protein SF3B1 is mutated. SF3B1 mutations are known to promote aberrant splicing and are associated with cancer, particularly the lysine 700 to glutamate mutation (K700E). We designed a pooled library of 249 *MAP3K7* mutants targeting branch points, RNA-binding protein motifs, nucleotide composition and predicted structural elements. The impact of these mutants on splicing was measured in the context of normal and SF3B1 K700E expression. RNA structure was assessed in parallel using *in vitro* high-throughput SHAPE-MAP chemical probing. We found that branchpoint mutations drive the strongest increases in cryptic splice-site use. There is no overall correlation between cryptic splice-site use and structural similarity to the wild-type *MAP3K7* RNA. However, mutants within an RNA binding protein hotspot (containing U2AF2, U2AF1, KHSRP and SRSF2 sites) are associated with cryptic splice-site use and structural similarity to wild-type *MAP3K7* RNA. These structural changes are associated with increased ensemble diversity. Our results demonstrate that although there are key structured regions within an RNA, there is also extensive variability where divergent RNA structures allow for accurate splicing.

## Introduction

Splicing factor 3b subunit 1 (SF3B1) protein is a core component of the spliceosome, with a role in branch point identification and 3’ splice site selection (Cretu et al. 2016; Tholen 2024). The *SF3B1 K700E* mutation, which is common in myelodysplastic syndromes, increases cryptic 3’ splice site use in a subset of genes (Darman et al. 2015; Dolatshad et al. 2015; Alsafadi et al. 2016; Damianov et al. 2025; Herbert et al. 2025). These cryptic sites activated by SF3B1 K700E mutant protein are typically expressed at low levels or remain silent in cells expressing wild-type SF3B1. In contrast, some genes contain cryptic splice sites that maintain similar levels of usage in both wild-type and SF3B1 K700E backgrounds. SF3B1 mutation-sensitive splice junctions share several recurrent features. Cryptic 3’ splice sites activated in SF3B1 K700E mutants are 10 to 30 base pairs upstream of the canonical splice site, are flanked in both directions by weak poly-pyrimidine tracts and frequently contain multiple branchpoints (Darman et al. 2015; Alsafadi et al. 2016; Herbert et al. 2025). Previously, we showed that SF3B1 mutation-sensitive junctions exhibit greater intronic ensemble diversity, meaning that these junctions have multiple predicted structures of similar energy (Herbert et al. 2025). However, the functional importance of ensemble diversity for accurate splice-site selection remains unclear.

To directly interrogate precursor RNA features that influence splicing accuracy, we employed a high-throughput structure:function assay centered on a *mitogen-activated protein kinase kinase kinase 7 (MAP3K7)* intron reporter. The *MAP3K7* gene is highly sensitive to mis-splicing in the context of SF3B1 K700E mutation and has been used in prior studies as a splicing minigene (North et al. 2022). We designed a pooled library of 249 mutants targeting branch points, polypyrimidine tracts, RNA-binding protein motifs, nucleotide composition, and predicted RNA structures. We assayed each mutant for its impact on splicing in normal SF3B1 and in the context of SF3B1 K700E expression. In parallel, we performed *in vitro* experimental chemical probing on these same mutants, enabling direct comparison of sequence, RNA structure and functional splicing outcomes. We found that a wide variety of divergently structured RNAs are still capable of accurate splicing. We also found that branchpoint motifs and RNA-binding protein sites strongly influence splice-site selection in the *MAP3K7* mini-gene in the context of both wild-type SF3B1 and SF3B1 K700E expression.

## Results

### High-throughput profiling of RNA structure and splicing in a pooled *MAP3K7* library

We designed a high-throughput method to capture splicing changes and RNA structure data from a single, multiplexed pool of 249 mutant RNAs (Figure 1A, Supplemental Table 1). We based our experiment on a previously developed SF3B1 K700E-sensitive mini-gene assay derived from the *MAP3K7* gene that includes the first 100 intronic bases from the 5’ splice site and last 150 intronic bases to the 3’ splice site (North et al. 2022). We modified the *MAP3K7* intron mini-gene so that the intron is flanked by partial segments of its endogenous *MAP3K7* exons and truncated the flanking GFP to shorten the construct (Figure 1A). We tested our truncated *MAP3K7* mini-gene by transfecting, extracting cDNA and amplifying across the exon junction. Use of the cryptic 3’ splice site in wild-type *MAP3K7* was increased when co-transfected with *SF3B1 K700E* (Figure 1B). The modified *MAP3K7* mini-gene was used as a template for the creation of a mutant library. Mutants were selected by targeting branchpoint motifs, GC content, structural elements and RNA binding protein sites based ENCODE data (ENCODE Project Consortium 2012). A major issue with high-throughput structure mapping of similar molecules is that sequencing reads cannot be accurately assigned to the correct construct. To overcome this, we used unique, hairpin-forming barcodes on each end of the construct based on recent high throughput structural assays (Figure 1A) (https://github.com/jyesselm/rna_lib_design.git) (Deenalattha et al. 2025). These barcodes are 25-mers with unique base-pairing 8-mers on each end and an identical internal 9-mer loop and were designed to enable the accurate retrieval of chemical probing reactivity data for each mutant RNA out of the multiplexed pool.

**Figure 1.**
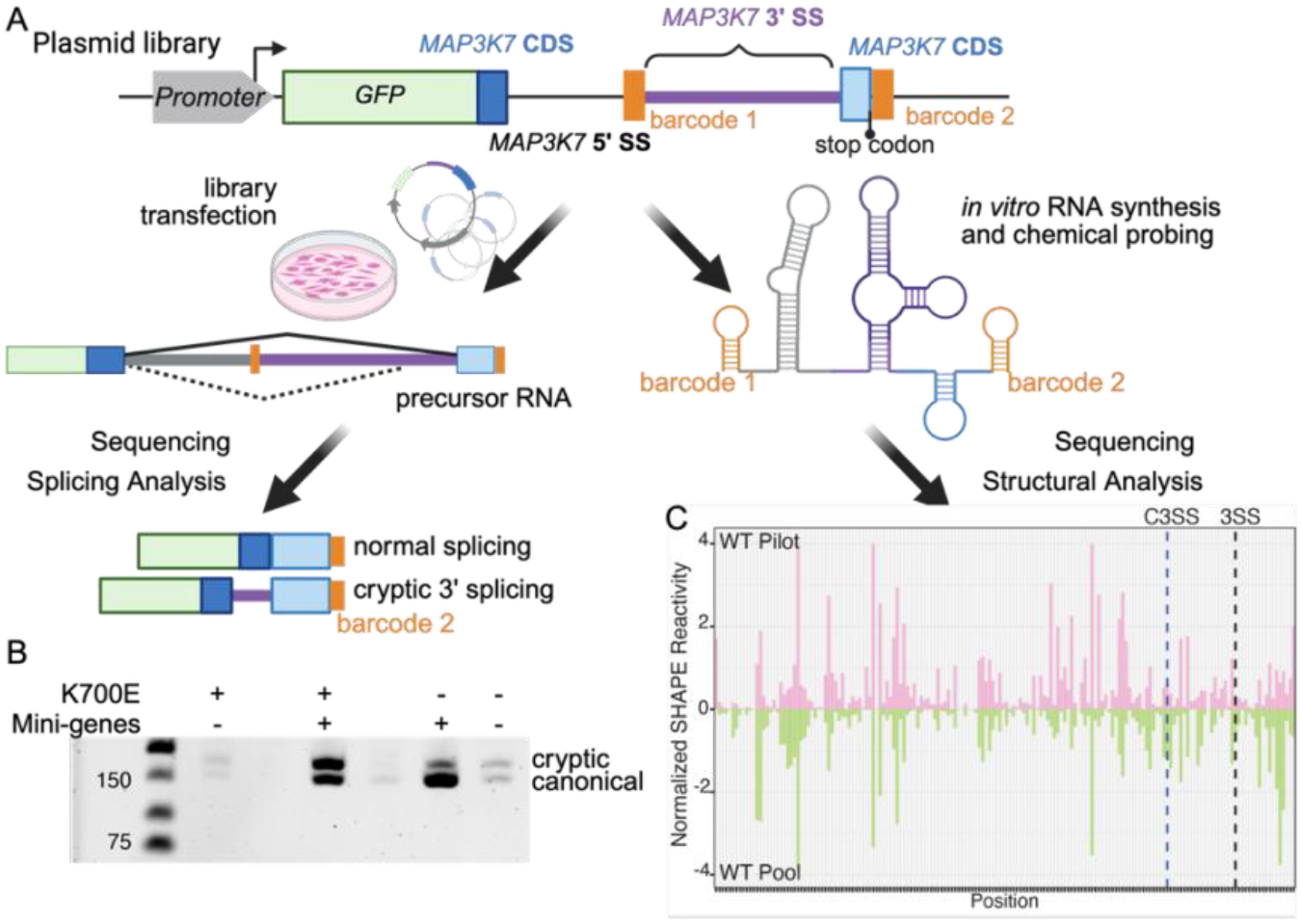
High-throughput structure-function mutagenesis screen for readouts on splicing and structure. (A) Synthetic intron construct designed for cellular transfection and alternative splicing analysis. Barcodes unique to each mutant oligo facilitate read demultiplexing. For splicing analysis, the mutant oligo pool was cloned into an expression vector, transfected into HEK293 cells, and sequenced. For RNA structure probing experiments the oligo pool was *in vitro* transcribed into RNA, structure probed using 5NIA, reverse transcribed into cDNA libraries and sequenced. (B) Cellular validation of the synthetic *MAP3K7* intron and its sensitivity to SF3B1 K700E mutation. (C) Normalized SHAPE reactivities for the wild type *MAP3K7* RNA structure probed in a small pilot (top, pink) versus as a part of the total mutant RNA pool (bottom, green) including the cryptic (C3SS) and canonical (3SS) 3’ splice sites. Created with BioRender.com.

As a small scale test, we pooled wild-type *MAP3K7*, a branch point mutant, and a structural mutant and performed selective 2′-hydroxyl acylation analyzed by primer extension and mutational profiling (SHAPE-MaP) (Siegfried et al. 2014). We compared the structural data from our pilot RNAs to structural data from the entire pool of wild-type and mutant *MAP3K7* RNAs. The wild-type *MAP3K7* from the pilot group has similar SHAPE reactivity to the wild-type *MAP3K7* from the complete pool (Spearman correlation = 0.61, *p* = 1.06e-19) (Figure 1C). We performed the same splicing and structural experiments on the full library of 249 mutants. We achieved high reproducibility between SHAPE replicates for each demultiplexed construct (median Spearman correlation = 0.76) (Figure 2A). To confirm that our barcode and demultiplexing step was segregating reads accurately, we created a baseline comparison by randomly sub-sampling reads from the entire pool and mapping these reads to the wild-type *MAP3K7*. Using five groups of randomly sub-sampled reads, we confirmed that the sub-sampled groups have much lower correlation to correctly demultiplexed *MAP3K7* constructs than when the pairs are properly matched (Figure 2B).

**Figure 2.**
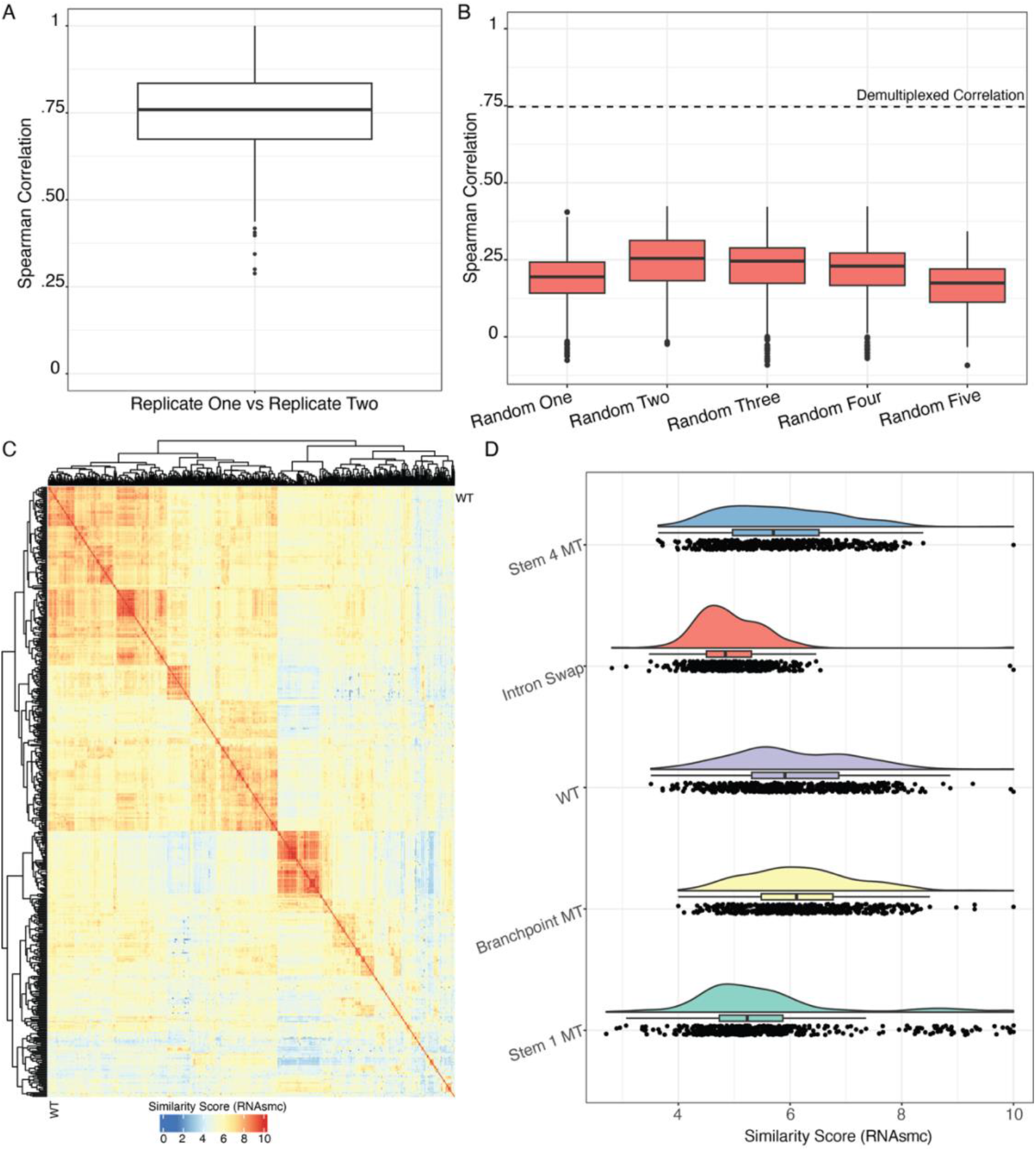
Per-mutant structure data is efficiently captured from pooled experiments. (A) Two replicates of chemical probing were demultiplexed and the SHAPE reactivity data for the matching samples were compared using Spearman correlation. (B) Correlation is not based on overall similarity as five randomly sub-sampled sets of reads mapped to the wild-type sequence exhibit low correlation in each instance. Median correlation between replicates from (A) plotted as a dashed line. (C) Hierarchical clustering of structural similarity scores (RNAsmc) shows a variety of structures among the entire pool of mutant RNAs. (D) Distribution of structure similarity scores (RNAsmc) for five selected RNAs shows different structural landscapes when compared against demultiplexed RNAs from the total pool.

We analyzed the distribution of structures across our pool. We took each *MAP3K7* RNA reactivity dataset and used it to create a minimum free energy model (MFE) representative structure. We compared the wild-type *MAP3K7* MFE to each mutant to derive a structural similarity score (RNAsmc) (Wang et al. 2023). Overall, our pool has a wide distribution of structures ranging from highly similar MFEs (above 9) to distinctly different MFEs (below 2) (Figure 2C). We found that the wild-type *MAP3K7* has a broad structural distribution (Figure 2D, WT), while more drastic mutants, such as a complete intron swap, have lower similarity to the majority of the *MAP3K7*-derived pool (Figure 2D, Intron Swap). Mutants with minimal changes and little expected impact on structure, such as the branchpoint mutant, showed a very similar distribution to the wild-type *MAP3K7* (Figure 2D, Branchpoint MT). This supports a wide range of structural arrangements within the pool.

### Structural similarity to wild-type *MAP3K7* does not predict cryptic splice-site use

To capture splicing data, we transfected the pool of *MAP3K7* constructs into HEK293 cells. We identified mutations that impact basal *MAP3K7* splicing by comparing the cryptic 3’ splice site use between each mutant RNA and the wild-type *MAP3K7* (ΔC3SS MT oligo - WT oligo). As expected, control mutants that disrupt splicing motifs significantly increase cryptic splice site use, including knocking out the canonical splice site or weakening the pyrimidine tract of the canonical splice site (Figure 3A, labeled). We then assessed RNA structure for the same *MAP3K7* pool with SHAPE-MaP chemical probing (Figure 1A). To analyze how each mutant RNA changed structure compared to the wild-type *MAP3K7*, we used normalized SHAPE reactivity data and performed Spearman correlations between each mutant and the wild-type in contrast to splicing accuracy (Figure 3B). Then, we compared experimentally informed mutant and wild-type MFE structures using RNAsmc to splicing accuracy (Wang et al. 2023) (Figure 3C). The change in cryptic 3’ splice site use was not significantly correlated with structural similarity using either metric (Figure 3B and C, Supplemental Table 4). We did find that both our methods of quantifying structural similarity were consistent, as there is a significant positive association between the correlation of normalized SHAPE reactivities and the comparison of MFE structures (Figure 3D, Supplemental Table 4), supporting the use of either raw experimental data correlations or minimum free energy model comparisons for structural analysis. Overall structural resemblance to wild-type MAP3K7 was insufficient to explain cryptic splice-site use.

**Figure 3.**
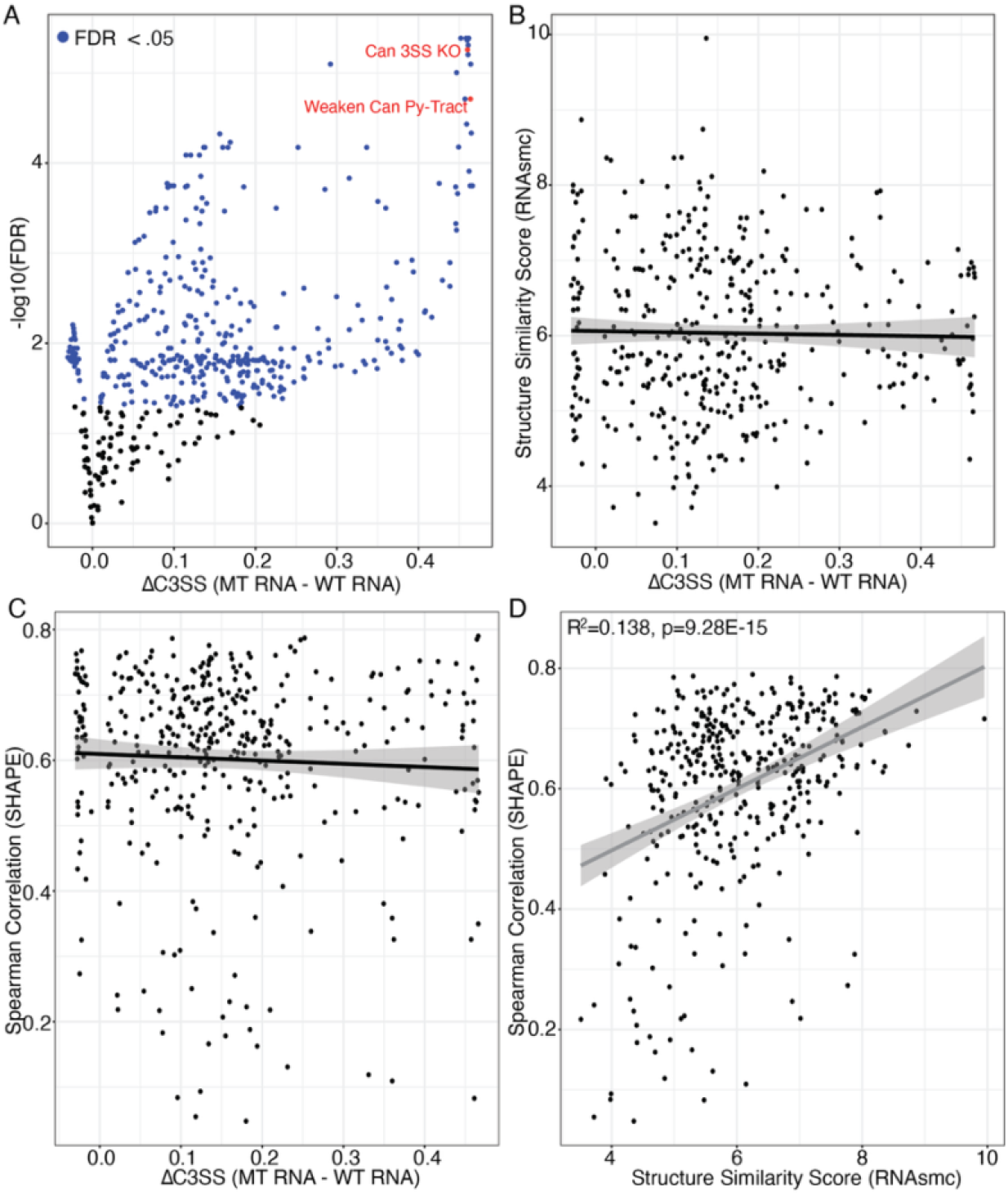
Splicing changes are not correlated with structural differences between mutant and wild-type *MAP3K7* RNAs. (A) Change in cryptic 3’ splice site use observed between mutant RNAs and wild-type *MAP3K7* RNA (FDR < .05, Welch’s two-sided t-test with Benjamini Hochberg correction). Two mutations designed to knock out the canonical 3’ splice site (Can 3SS KO) and disrupt the polypyrimidine tract (Weaken Can Py-Tract) show strong increases in cryptic 3’ splice site use (indicated in red) as expected. (B) Comparison of RNA similarity scores between mutant and wild-type *MAP3K7* (RNAsmc) shows no significant correlation with change in cryptic 3’ splice site use. (C) Comparison of normalized SHAPE reactivities between mutant and wild-type *MAP3K7* show no significant correlation with change in cryptic 3’ splice site use. (D) Structure method comparison of similarity scores (RNAsmc) and normalized SHAPE reactivities between each mutant and wild-type *MAP3K7* shows significant positive correlation. Statistics in Supplemental Table 4.

Second, we assessed the effect of expression of SF3B1 K700E on cryptic splice site usage for each *MAP3K7* RNA in our pool. This comparison identifies mutants with exacerbated cryptic 3’ splicing when SF3B1 K700E is expressed (ΔC3SS K700E-WT). We observed a significant change in cryptic splice site use in many mutant *MAP3K7* RNAs with expression of SF3B1 K700E, including a proof-of-concept RNA with a strengthened polypyrimidine tract at the cryptic 3’ splice site that increases cryptic splice site use as expected (Figure 4A, labeled). Comparing the change in cryptic splice site use to normalized SHAPE reactivity data correlations between each mutant RNA and wild-type *MAP3K7* revealed no significant association (Figure 4B, Supplemental Table 4). Likewise, comparing the change in cryptic splice site use to the MFE-based structural similarity (RNAsmc) yielded no significant relationship (Figure 4B, Supplemental Table 4). These data indicate that splicing accuracy is compatible with a broad range of RNA structural conformations.

**Figure 4.**
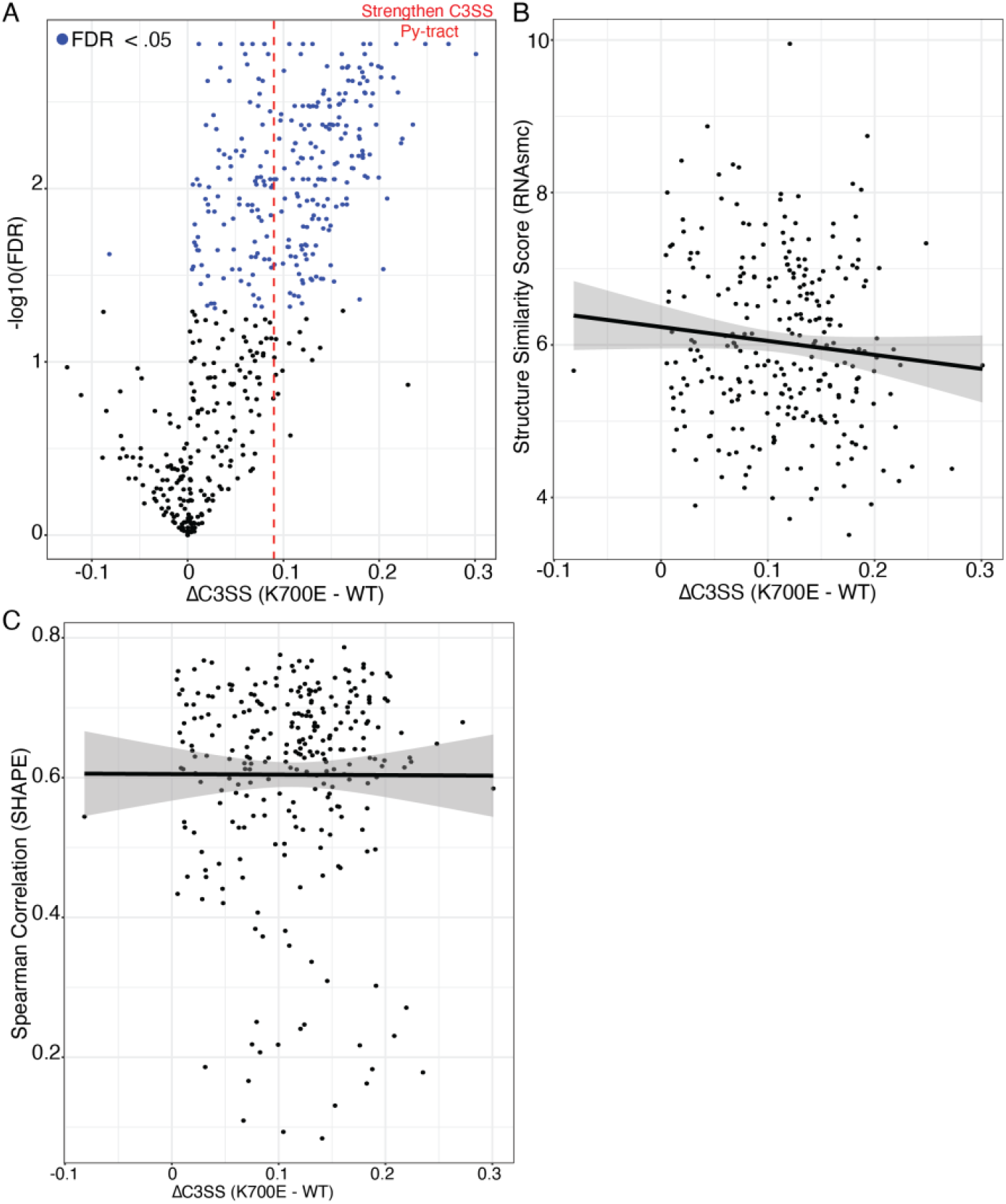
SF3B1 K700E induced cryptic 3’ splice site changes are not correlated with structural differences between mutant and wild-type *MAP3K7* RNAs. (A) Co-transfection of a *SF3B1 K700E* construct changes to cryptic 3’ splice site use. Splicing changes in the wild-type *MAP3K7* are indicated by a dashed red line. We confirmed that a mutant expected to increase cryptic splice site use by strengthening the cryptic polypyrimidine tract shows higher levels of cryptic splicing (Strengthen C3SS Py-Tract). (B) Comparison of structural similarity scores between mutant and wild-type *MAP3K7* (RNAsmc) with SF3B1 K700E-induced splicing changes shows no significant correlation. (C) Comparison of normalized SHAPE reactivities between mutant and wild-type *MAP3K7* with SF3B1 K700E-induced cryptic splicing changes shows no significant correlation. Statistics in Supplemental Table 4.

### Different subclasses of *MAP3K7* mutants show distinct relationships with RNA structure

To determine how intronic regulatory elements differentially contribute to cryptic 3′ splice site activation, we analyzed mutant *MAP3K7* RNAs by category. Categories were based on overlap of mutations with potential functionality in the wild-type *MAP3K7* intron. Mutation categories with the largest enrichment of significant splicing changes were selected for further investigation, including branchpoint motifs (yUnAy), RNA binding protein (RBP) Block Three (U2AF2, SRSF2, KHSRP, and U2AF1 binding sites) and RBP Block One (SRSF1, SRSF2, and BUD13 binding sites) (Figure 5A).

**Figure 5.**
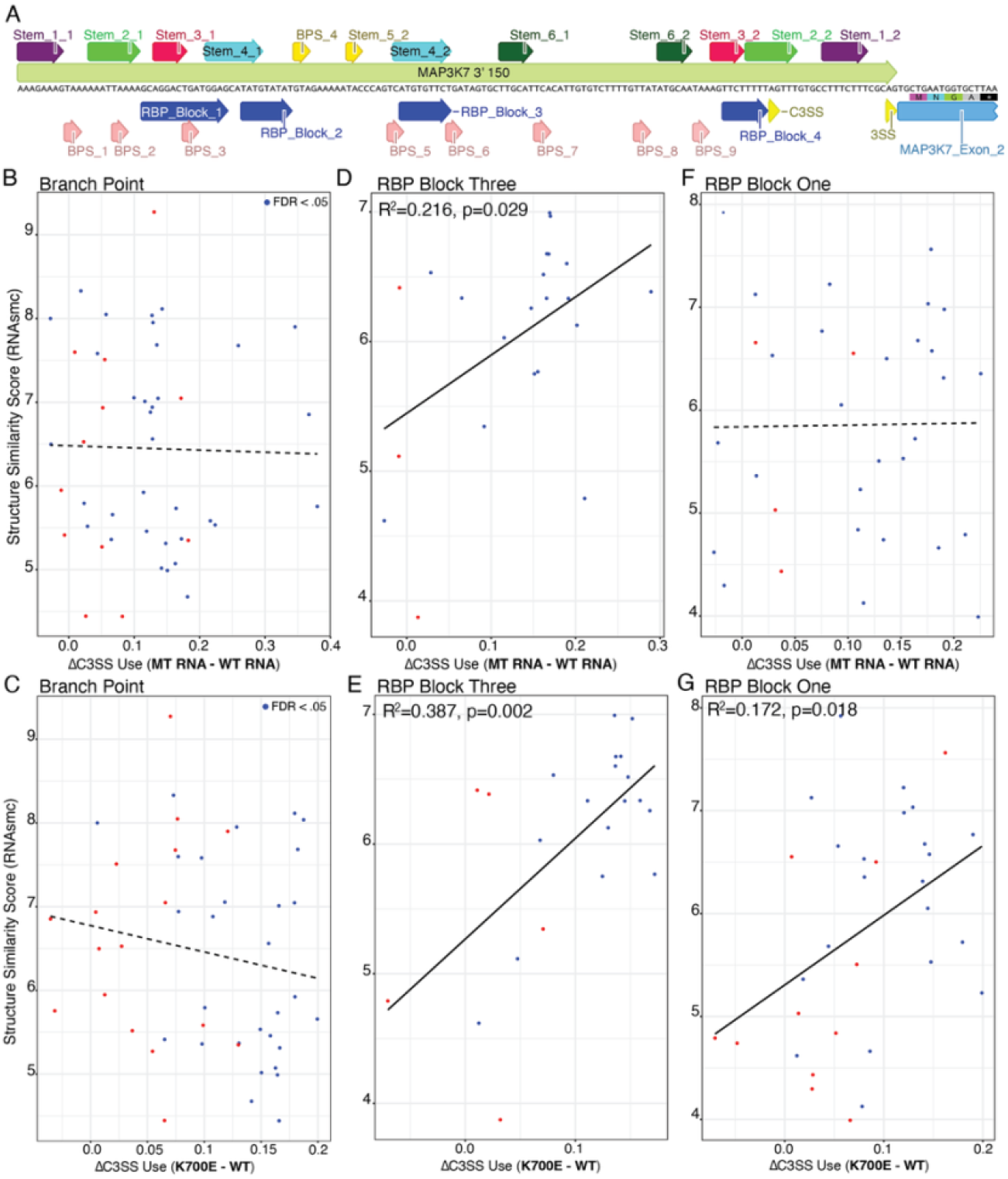
Mutation within branchpoint, RBP Block Three, or RBP Block One regions significantly alters cryptic 3’ splice site use. (A) Gene map of annotations on the wild-type *MAP3K7* synthetic RNA. (B) Branchpoint mutants show no significant correlation between cryptic 3’ splice site use and RNAsmc structural similarity scores in normal SF3B1 background or (C) with SF3B1 K700E expression. (D) RBP Block Three mutant splicing changes correlated positively with structural similarity to wild-type *MAP3K7* and (E) were exacerbated by SF3B1 K700E expression. (F) RBP Block One mutant splicing changes show no correlation with structural similarity to wild-type *MAP3K7* in normal cells, but (G) expression of SF3B1 K700E results in significant correlation between splicing and structural similarity to wild-type *MAP3K7*. Statistics in Supplemental Table 4.

Mutant RNAs with a significant increase in cryptic splice site use were enriched for branchpoint mutations. This occurred at the basal splicing level, with 37 out of 46 branchpoint mutant RNAs showing significant baseline splicing changes (mean increase in cryptic splice site use = 0.137, Supplemental Tables 2 and 3). In addition, SF3B1 K700E expression exacerbated cryptic splice use, with 32 out of 46 branchpoint mutant RNAs showing significant increases (mean increase in cryptic splice site use = 0.136, Supplemental Table 5 and 6). However, there is no correlation between impact on cryptic splice site use and structural similarity within the branchpoint mutant pool in either the wild-type SF3B1 context or with expression of SF3B1 K700E (Figure 5B and C, Supplemental Table 4), similar to our observations across the total *MAP3K7* pool (Figure 3B and C). As this category selects mutants that specifically change the branchpoint sequence motif, it is not surprising to find a lack of correlation between splicing and RNA structure. Interestingly, the expression of SF3B1 K700E does alter cryptic splice site use, supporting an important role for branchpoint integrity in cryptic splice site activation.

*MAP3K7-*derived RNAs with mutations overlapping the RBP Block Three–associated elements, including Stem 4, showed greater structural dependence. RBP Block Three contains RNA binding protein interaction sites for U2AF2, SRSF2, KHSRP, and U2AF1, as identified through analysis of ENCODE eCLIP data (ENCODE Project Consortium 2012). 19 out of 22 RNAs with mutations in RBP Block Three had significant changes in cryptic splice site use in the SF3B1 wild-type background (mean increase in cryptic splice site use = 0.148, Supplemental Table 2 and 3). Furthermore, 19 out of 22 mutant RNAs also had a significant increase in cryptic splice site use with the SF3B1 K700E mutation (mean increase in cryptic splice site use = 0.122, Supplemental Tables 5 and 6). Interestingly, RBP Block Three mutants show a significant positive correlation between structure similarity to the wild-type *MAP3K7* RNA and both basal and SF3B1-mediated cryptic 3’ splice site use (Figure 5D and E, Supplemental Table 4). This is surprising, because it means that mutants that are more structurally similar to wild-type have increased cryptic splice site use. These data suggest that the structural similarity of RBP Block Three to wild-type *MAP3K7* promotes the cryptic splice site and disruption of RBP Block Three structure may allow for more canonical splicing.

Finally, mutants overlapping RBP Block One, which includes Stem 3, were also enriched in increased cryptic splicing. RBP Block One contains RNA binding protein motifs for SRSF1, SRSF2, and BUD13 (ENCODE Project Consortium 2012). 28 out of 32 constructs with mutation within the RBP Block One region showed significant change in cryptic 3’ splice site values (mean increase in cryptic splice site use = 0.114, Supplemental Table 2 and 3). In addition, in the context of SF3B1 K700E expression, 23 out of 32 constructs with mutation within RBP Block One had significant increase in cryptic 3’ splice site use (mean increase in cryptic splice site use = 0.104, Supplemental Table 5 and 6). The splicing perturbation of this group of mutant RNAs is not significantly correlated with changes in RNA structure under normal conditions (Figure 5F, Supplemental Table 4). However, with SF3B1 K700E expression, the splicing changes by mutants in RBP Block One are associated with structural similarity to wild-type *MAP3K7* (Figure 5G, Supplemental Table 4). This is similar to RBP Block Three, where structure similarity to wild-type in this region seems to promote cryptic splice site selection. These findings suggest that RBP Block One–associated elements contribute strongly to both intrinsic and SF3B1 K700E-mediated splice site selection, but structural similarity to wild-type *MAP3K7* RNA most important in the context of SF3B1 K700E expression.

### RBP-associated MAP3K7 mutants exhibit increased ensemble diversity and cryptic splice-site usage

RNAs commonly exist in an ensemble of different structures. Some RNAs have one or two structures with low energy that are likely to dominant the ensemble, while other RNAs can form many structures that have similar energy. To analyze the ensemble diversity and structural stability of our pool of *MAP3K7-*derived mutant RNAs, we used Scanfold (Andrews et al. 2018). Ensemble diversity measures variation in base-pairing across Boltzmann-weighted RNA conformations, with higher values indicating increased structural flexibility (Andrews et al. 2018; Moss 2018). In addition, Scanfold also detect stable structures by using the MFE structure as a baseline to compare shuffled versions of the same sequence, with more negative Z scores indicating more stable folds (Andrews et al. 2018). We compared the ensemble diversity and structural stability of constructs with mutations in the branchpoint, RBP Block Three and RBP Block One to the wild-type *MAP3K7* RNA and remaining additional mutants. Analysis of the mean per-base ensemble diversity and structural stability revealed overall more ensemble diversity and less stable structures for RBP Block One and RBP Block Three mutants (Figure 6A and B). In contrast, the branchpoint mutant RNAs had lower structural stability, similar to wild-type *MAP3K7* (Figure 6C). This data suggests that mutants in RBP Block One and Three influence RNA flexibility.

**Figure 6.**
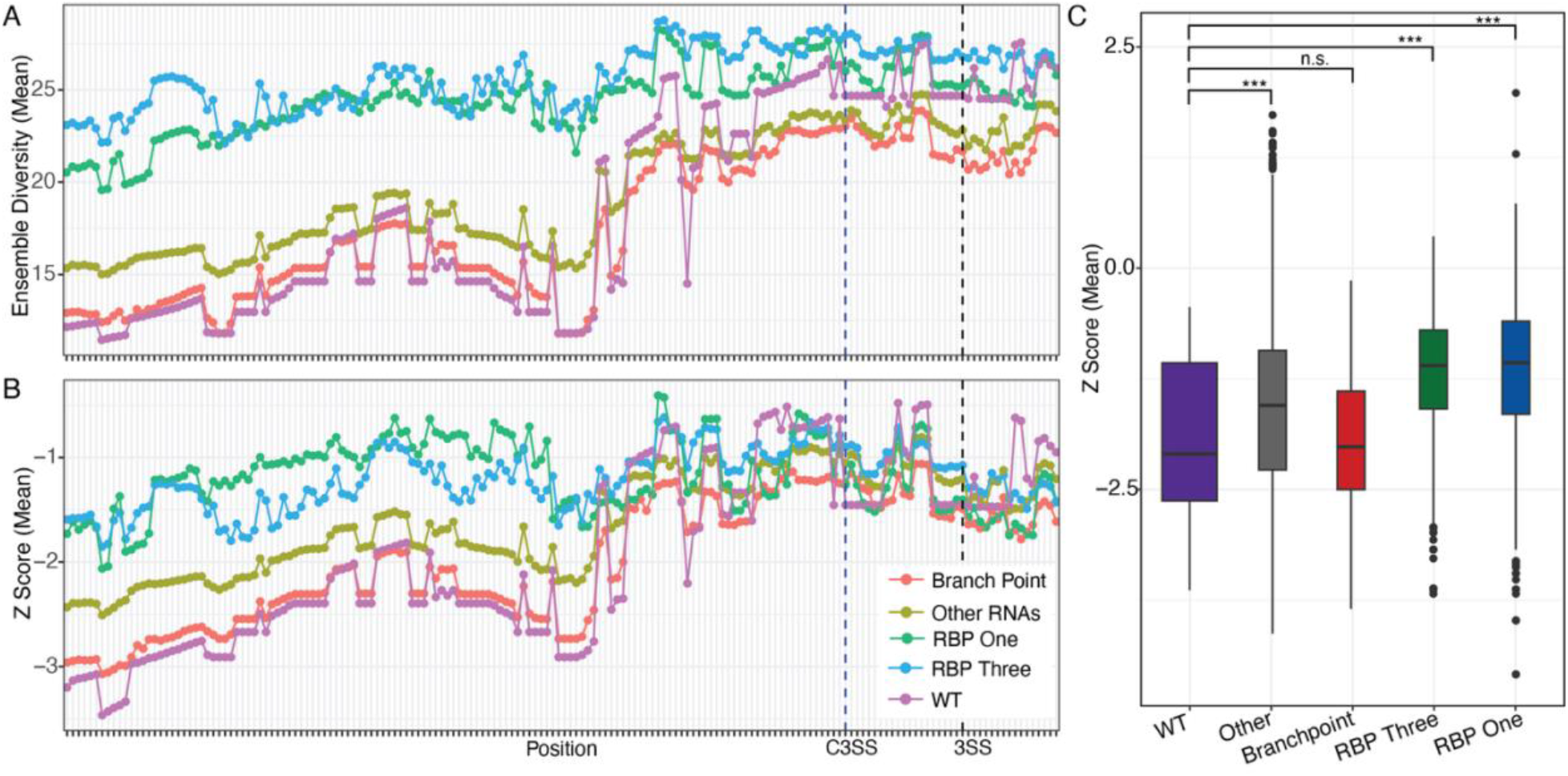
RBP Block One and Three mutants have higher ensemble diversity and fewer stable structures than the majority of *MAP3K7-*derived RNAs. (A) Per-base mean ensemble diversity for each category of RNAs. Branchpoint mutants (red), all other mutants not including the plotted groups (brown), RBP Block One (green), RBP Block Three (blue) and wild-type *MAP3K7* (purple). (B) Per-base average Z scores for each category of RNAs. Same colors as (A). (C) Box plot of mean Z scores for each gene across each mutant in a category. *** p-value < 0.001.

## Discussion

High-throughput structure:function assays permit cost-effective readouts on both splicing changes and *in vitro* structure differences for many individual mutated RNAs within a pool. By assaying hundreds of defined intronic perturbations within a single synthetic MAP3K7 intron, we can measure how sequence and RNA secondary structure contribute to splice-site choice. We can also add perturbations, like SF3B1 K700E expression. Interestingly, across the full *MAP3K7-*derived mutant pool we find no relationship between changes in cryptic splice-site usage and similarity to wild-type *MAP3K7* RNA structure (Figure 3B and C). Correspondingly, structural similarity to wild-type *MAP3K7* RNA structure does not correlate with splicing sensitivity to SF3B1 K700E mutation (Figure 4B and C). We find that *MAP3K7* RNA structures are highly malleable, and many diverse structural conformations remain capable of accurate splice site selection and completion of canonical splicing.

Branchpoint mutations produce the strongest enhancement of cryptic 3′ splice site usage. Disruption of splicing by branchpoint mutation is not correlated with structural similarity to *MAP3K7* (Figure 5B and C). The importance of branchpoint sequences throughout the intron support a key role for branchpoint integrity in SF3B1 mutation sensitivity and are consistent with models in which SF3B1 mutations alter the interpretation of branch point sequence or positioning during early spliceosome assembly (Darman et al. 2015; Alsafadi et al. 2016; Damianov et al. 2025).

In contrast, mutations overlapping RBP Block Three–associated elements exhibit a significant positive correlation between structural similarity and splicing perturbation (Figure 5D and E). This region contains predicted eCLIP binding sites for U2AF2, U2AF1, KHSRP, and SRSF2, all of which have established roles in early 3′ splice site recognition or splice-site selection (ENCODE Project Consortium 2012; Yan et al. 2019; Van Nostrand et al. 2020; Tholen 2024). We also found that RBP Block Three mutants had higher ensemble diversity and fewer local stable structures, suggesting that wild-type structures here are important for accessibility. Mutants in RBP Block One only show correlation between splicing efficiency and structural similarity to wild-type *MAP3K7* in the context of SF3B1 K700E expression (Figure 5F and G). This region contains predicted binding sites for SRSF1, SRSF2, and BUD13, factors that are implicated in exon definition, splice-site strengthening, and spliceosome assembly (Long and Caceres 2009; Zhou and Fu 2013; Bertram et al. 2017). Like in RBP Block Three, this correlation is positive, suggesting that the cryptic splice site is enhanced by RNA structures similar to wild-type *MAP3K7* in this region. RBP Block One mutants that increase in cryptic splice site use in the context of SF3B1 K700E expression also have higher ensemble diversity and fewer stable local structures, suggesting that this region may influence splice-site competitiveness.

## Materials and Methods

### Cell Culture

HEK293 cells (ATCC) were maintained in DMEM medium (Gibco) supplemented with 10% fetal bovine serum (FBS), 1% penicillin-streptomycin at 37°C with 5% CO2 and passaged every three days. Cells were checked regularly for mycoplasma contamination.

### Oligo design

A SF3B1-sensitive *MAP3K7* intron mini-gene has been previously tested and reported (North et al. 2022). Based on the previous construct, we designed a gene fragment of *MAP3K7* composed of the first 100 base pairs of the intron from the 5’ splice site and the last 150 base pairs leading to the 3’ splice site. The wild-type, branch point mutant and stem structure mutants were ordered as gene fragments from Integrated DNA Technologies (IDT), PCR amplified using Q5 TAQ (New England Biolabs, NEB) and cloned into a pcDNA3.1 backbone using a Hifi cloning kit (NEB) (Primers: LL583, LL579). These three test constructs were transfected into HEK293 cells with or without co-transfection of a SF3B1 K700E expression plasmid using the Lipofectamine 3000 transfection kit (ThermoFisher). Transfected cells were incubated at 37 C for two days and whole-cell RNA was purified using a standard Trizol procedure followed by cleanup with the Monarch Spin RNA isolation kit (NEB). RNA was reverse transcribed using gene specific primer with the SuperScript IV RT kit (ThermoFisher) and cDNA was subsequently amplified using the Q5 TAQ (NEB) under standard PCR conditions. PCR products were run out on a 6% TBE gel (Invitrogen) and stained using SYBRsafe gel dye (Invitrogen) for one hour at room temperature. All RNAs designed in this study are available in Supplemental Table 1. All primers utilized in this study are available in Supplemental Table 7.

### Barcode oligos

Mutant oligos with unique hairpin forming barcodes and homology arms on each end were ordered and synthesized as gene fragments (IDT). Unique barcodes added to each end enable accurate demultiplexing of reads by barcode and by forming a hairpin to themselves, prevent SHAPE modification during structure probing as demonstrated in previous work (Deenalattha et al. 2025). Unique barcodes were designed as 25-mers with an 8-mer on each end forming the stem and the same internal 9-mer that forms the bulge in the hairpin. Barcode information for each RNA is available in the sequences in Supplemental Table 1.

### In vitro SHAPE probing

The synthesized oligo pool was PCR amplified using the Q5 TAQ (New England Biolabs, NEB) with a generic reverse primer and forward primer containing a T7 promotor (Supplemental Table 7: LL582F, LL583R). Using the PCR products as template, *in vitro* RNA synthesis was performed for two hours at 37°C using the HiScribe T7 High Yield RNA Synthesis Kit (New England Biolabs). *In vitro* synthesized RNA was DNA digested with TurboDNAse and purified with RNA XP cleanup beads (Beckman Coulter). RNA was incubated in folding buffer (400 mM bicine, 200 mM NaCl, 20 mM MgCL2) and 25 mM 5NIA or DMSO at 37°C for 5 minutes and immediately cleaned up using RNA XP beads (Beckman Coulter) and input into error-prone reverse transcription using the Invitrogen Super Script II (ThermoFisher Scientific) with modified RT buffer (10X: mM Tris pH 8, 750 mM KCl) supplemented with 6 nM manganese, RNAse inhibitor, and a gene specific primer (Supplemental Table 7, LL583). The RT product was cleaned up with the RNA XP beads and used as template for a PCR reaction of 12 cycles using template specific primers (Supplemental Table 7: LL588, LL589). Products from the first PCR were cleaned up using RNA XP beads and used as template for a second PCR reaction of 10 cycles using Tru-Seq uniquely barcoded primers. Libraries were cleaned up using the RNA XP kit and sequenced and sequenced (2 ×150, Illumina).

### Analysis of SHAPE data

SHAPE analysis was conducted on the raw sequencing reads using Shapemapper-txome v1.0 under default parameters with SHAPE treated reps as modified and DMSO treatments as untreated. SHAPE reactivities determined from Shapemapper-txome were used to guide prediction of minimum free energy (MFE) structures for each structure using the ViennaRNA v2.4.18 RNAfold command under default parameters (Lorenz et al. 2011; Busan and Weeks 2018). Dot bracket MFE structures were used to determine structural similarity scores using RNAsmc v0.8.0 under default parameters and visualized in R using pheatmap v1.0.13 package (Wang et al. 2023). Spearman correlation between RNAs normalized SHAPE reactivities was determined in R v4.5.2. Scanfold2 v1.0 with a step size of one and window size of 120 (Andrews et al. 2018). Per base ensemble diversities and Z scores were imported and analyzed in R.

### Splicing assay

The mutant, barcoded oligo pool was PCR amplified using generic primers with homology arms to the backbone (Supplemental Table 7, LL583, LL586) via the Q5 TAQ (New England Biolabs, NEB). The backbone was amplified using the same kit with generic primers matching the homology arms added to the oligos. Cloning was conducted using the NEBuilder Hifi DNA Assembly Cloning Kit (New England Biolabs). Colonies were incubated at 37 C overnight on Luria-Bertani agar supplemented with ampicillin (ThermoFisher). Cloning efficiency was determined by colony PCR of 10 colonies from each plate. Colony plates were scraped and used as starter for Luria-Bertani broth supplemented with ampicillin liquid cultures. Liquid culture was grown overnight at 37 C and plasmid was purified using the ZymoPURE II Maxiprep Kit (Zymo Research). These pooled plasmids were individually transfected into HEK293 cells with or without co-transfection of a SF3B1 K700E expression plasmid using the Lipofectamine 3000 transfection kit (ThermoFisher). Transfected cells were incubated at 37 C for two days and whole-cell RNA was purified using a standard Trizol procedure followed by cleanup with the Monarch Spin RNA isolation kit (NEB). RNA was reverse transcribed using gene specific primer (Supplemental Table 7, LL583) with the SuperScript IV RT kit (ThermoFisher) and cDNA was subsequently amplified using the Q5 TAQ (NEB) under standard PCR conditions for 8 cycles (LL589, LL596). First phase libraries were cleaned up using the RNA XP beads and underwent a second PCR reaction of 8 cycles using Tru-Seq primers. Final libraries were cleaned up using the RNA XP beads and sequenced (Illumina, 2×150). Each experiment was repeated in three replicates for SF3B1 K700E co-transfection and three paired controls.

### Splicing analysis

The raw reverse reads were demultiplexed by unique barcode using Shapemapper-txome v1.0 under default parameters using reads from the SF3B1 K700E co-transfection as the modified sample and SF3B1 WT reads as the untreated sample (Busan and Weeks 2018). Each paired replicate was processed individually. Demultiplexed reads were mapped back to the reference sequences using split-read mapping with STAR v2.7.10 using parameters “--outFilterMultimapNmax 1 --outFileNamePrefix --outSAMtype BAM SortedByCoordinate -- outSAMunmapped None --outSAMattributes Standard” (Dobin et al. 2013). Bam files were indexed using Samtools v1.10 index command with default setting. Per-nucleotide read counts for each RNA were determined with Samtools v1.10 using the Samtools depth command with parameters “-H -aa -d 0” (Li et al. 2009). Raw nucleotide counts were imported into R and transformed into counts per million. Then, two metrics were determined: splicing change for each mutant RNA versus the wild-type RNA under SF3B1 wild-type conditions and splicing change for each mutant RNA versus itself between SF3B1 K700E and SF3B1 WT conditions. Equations:

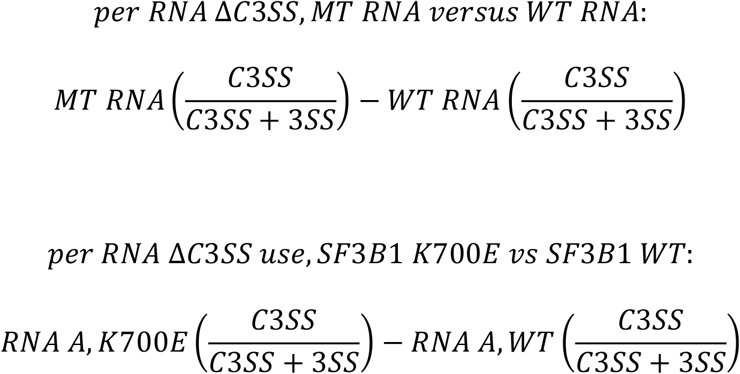

Data was visualized in R v4.5.2. Scripts available at: https://github.com/herber4/minigene_analysis_snakemake/tree/main

## Acknowledgments

We thank the Genomics and Bioinformatics Core of the Clemson University Center for Human Genetics for analytical and computational assistance with Secretariat High Performance computing cluster used for data analysis (supported by NIH grant P20GM139769).

## Author Contributions

Austin Herbert and Lela Lackey designed the experiments, coordinated the project and supervised the work. Austin Herbert, Alexandra Randazza and Abigail Hatfield performed the experiments. Austin Herbert performed data curation and data analysis. Austin Herbert and Lela Lackey wrote the original draft. Austin Herbert and Lela Lackey contributed to editing.

## Supplemental Tables

**Supplemental Table 1**. *MAP3K7* pool sequences.

**Supplemental Table 2**. Statistical analysis of MAP3K7 mutant categories for splicing accuracy and similarity to wild-type structure.

**Supplemental Table 3**. Individual values for MAP3K7 mutants for splicing accuracy (MT RNA-WT RNA) and similarity to wild-type structure.

**Supplemental Table 4**. Regression statistics for structure:function comparisons.

**Supplemental Table 5**. Statistical analysis of MAP3K7 mutant categories for SF3B1 K700E-mediated splicing accuracy and similarity to wild-type structure.

**Supplemental Table 6**. Individual values for MAP3K7 mutants for splicing accuracy (SF3B1 K700E - SF3B1 WT) and similarity to wild-type structure.

**Supplemental Table 7**. Additional primer sequences.

## References

Alsafadi S, Houy A, Battistella A, Popova T, Wassef M, Henry E, Tirode F, Constantinou A, Piperno-Neumann S, Roman-Roman S et al. 2016. Cancer-associated SF3B1 mutations affect alternative splicing by promoting alternative branchpoint usage. Nature Communications 7: 10615.

Andrews RJ, Roche J, Moss WN. 2018. ScanFold: an approach for genome-wide discovery of local RNA structural elements-applications to Zika virus and HIV. PeerJ 6: e6136.

Bertram K, Agafonov DE, Liu WT, Dybkov O, Will CL, Hartmuth K, Urlaub H, Kastner B, Stark H, Luhrmann R. 2017. Cryo-EM structure of a human spliceosome activated for step 2 of splicing. Nature 542: 318–323.

Busan S, Weeks KM. 2018. Accurate detection of chemical modifications in RNA by mutational profiling (MaP) with ShapeMapper 2. RNA 24: 143–148.

Cretu C, Schmitzova J, Ponce-Salvatierra A, Dybkov O, De Laurentiis EI, Sharma K, Will CL, Urlaub H, Luhrmann R, Pena V. 2016. Molecular Architecture of SF3b and Structural Consequences of Its Cancer-Related Mutations. Mol Cell 64: 307–319.

Damianov A, Lin CH, Zhang J, Manley JL, Black DL. 2025. Cancer-associated SF3B1 mutation K700E causes widespread changes in U2/branchpoint recognition without altering splicing. Proc Natl Acad Sci U S A 122: e2423776122.

Darman RB, Seiler M, Agrawal AA, Lim KH, Peng S, Aird D, Bailey SL, Bhavsar EB, Chan B, Colla S et al. 2015. Cancer-Associated SF3B1 Hotspot Mutations Induce Cryptic 3’ Splice Site Selection through Use of a Different Branch Point. Cell Rep 13: 1033–1045.

Deenalattha DHS, Jurich CP, Lange B, Armstrong D, Nein K, Yesselman JD. 2025. Characterizing 3D RNA structural features from DMS reactivity. bioRxiv.

Dobin A, Davis CA, Schlesinger F, Drenkow J, Zaleski C, Jha S, Batut P, Chaisson M, Gingeras TR. 2013. STAR: ultrafast universal RNA-seq aligner. Bioinformatics (Oxford, England) 29: 15–21.

Dolatshad H, Pellagatti A, Fernandez-Mercado M, Yip BH, Malcovati L, Attwood M, Przychodzen B, Sahgal N, Kanapin AA, Lockstone H et al. 2015. Disruption of SF3B1 results in deregulated expression and splicing of key genes and pathways in myelodysplastic syndrome hematopoietic stem and progenitor cells. Leukemia 29: 1092–1103.

ENCODE Project Consortium. 2012. An integrated encyclopedia of DNA elements in the human genome. Nature 489: 57–74.

Herbert A, Hatfield A, Randazza A, Miyamoto V, Palmer K, Lackey L. 2025. Precursor RNA structural patterns at SF3B1 mutation sensitive cryptic 3’ splice sites. RNA Biol 22: 1–15.

Li H, Handsaker B, Wysoker A, Fennell T, Ruan J, Homer N, Marth G, Abecasis G, Durbin R, Genome Project Data Processing S. 2009. The Sequence Alignment/Map format and SAMtools. Bioinformatics 25: 2078–2079.

Long JC, Caceres JF. 2009. The SR protein family of splicing factors: master regulators of gene expression. Biochem J 417: 15–27.

Lorenz R, Bernhart SH, Honer Zu Siederdissen C, Tafer H, Flamm C, Stadler PF, Hofacker IL. 2011. ViennaRNA Package 2.0. Algorithms Mol Biol 6: 26.

Moss WN. 2018. The ensemble diversity of non-coding RNA structure is lower than random sequence. Noncoding RNA Res 3: 100–107.

North K, Benbarche S, Liu B, Pangallo J, Chen S, Stahl M, Bewersdorf JP, Stanley RF, Erickson C, Cho H et al. 2022. Synthetic introns enable splicing factor mutation-dependent targeting of cancer cells. Nature Biotechnology 40: 1103–1113.

Siegfried NA, Busan S, Rice GM, Nelson JA, Weeks KM. 2014. RNA motif discovery by SHAPE and mutational profiling (SHAPE-MaP). Nat Methods 11: 959–965.

Tholen J. 2024. Branch site recognition by the spliceosome. RNA 30: 1397–1407.

Van Nostrand EL, Freese P, Pratt GA, Wang X, Wei X, Xiao R, Blue SM, Chen JY, Cody NAL, Dominguez D et al. 2020. A large-scale binding and functional map of human RNA-binding proteins. Nature 583: 711–719.

Wang H, Lu X, Zheng H, Wang W, Zhang G, Wang S, Lin P, Zhuang Y, Chen C, Chen Q et al. 2023. RNAsmc: A integrated tool for comparing RNA secondary structure and evaluating allosteric effects. Comput Struct Biotechnol J 21: 965–973.

Yan C, Wan R, Shi Y. 2019. Molecular Mechanisms of pre-mRNA Splicing through Structural Biology of the Spliceosome. Cold Spring Harb Perspect Biol 11.

Zhou Z, Fu XD. 2013. Regulation of splicing by SR proteins and SR protein-specific kinases. Chromosoma 122: 191–207.

